# Long-term stability of dominance hierarchies in a wild parrot with fission-fusion dynamics

**DOI:** 10.1101/2024.05.15.594429

**Authors:** Julia Penndorf, Damien Farine, John Martin, Lucy Aplin

**Affiliations:** Cognitive and Cultural Ecology Research Group, Max Planck Institute of Animal Behavior, Radolfzell, Germany; Division of Ecology and Evolution, Research School of Biology, Australian National University, Canberra, ACT, Australia; Department of Evolutionary Biology and Environmental Studies, University of Zurich, Zurich, Switzerland; Department of Collective Behavior, Max Planck Institute of Animal Behavior, Konstanz, Germany; Hawkesbury Institute for the Environment, Western Sydney University, NSW, Australia

## Abstract

Dominance hierarchies are a common feature of stable groups, allowing animals to limit the costs of fighting over access to resources. However, while the emergence of dominance is relatively well known from species that form stable groups, less is known about whether hierarchies are maintained in societies with open group membership. One challenge for species that live in an open social environment is that they have the possibility of engaging in dominance interactions with a large number of conspecifics. To better understand how animals navigate the complexities of interacting within a large, open society, we recorded social associations and aggressive interactions in a highly social, communally roosting parrot, the sulphur-crested cockatoo (*Cacatua galerita*). By following 515 individuals across three neighbouring communities and recording social interactions during foraging, we show that sulphur-crested cockatoos form clear, linear hierarchies. We find males rank higher than females and that adults rank higher than juveniles. Within sex, where individuals may still have to compete with up to 100 individuals from the same age-sex class, body size did not affect dominance rank. Finally, we find that, despite highly dynamic social associations (fission-fusion dynamics) among individuals, hierarchies are stable, with dominance ranks being highly repeatable across at least three years. This study demonstrates that closed group membership is not a pre-requisites for stable dominance hierarchies to emerge.

## Introduction

Contests over limited resources are a fact of life for most social species (Ward and Webster, 2016), and can potentially be costly for the animals involved. As a result, many animals have evolved mechanisms to assess competitors, allowing them to target a subset of potential competitors to engage with and decrease overall level of aggression (Drews, 1993; Arnott and Elwood, 2009; Hobson and DeDeo, 2015; Hobson et al., 2021; Dehnen, Papageorgiou, et al., 2022). Dominance hierarchies are one such mechanism, and dominance rank is thought to give the most precise information on competitive ability (Parker, 1974). Accordingly, the question of how animal groups form and maintain dominance hierarchies has received much attention over the last century (Dehnen, Arbon, et al., 2022; Hobson, 2022).

Current research about the formation and maintenance of dominance hierarchies focuses around two axes. The first considers how and when contests are determined by the intrinsic resource holding potentials (RHP) of individuals. RHP is typically based on a combination of morphological (e.g., body-size or colouration Arnott and Elwood, 2009) and behavioural traits (e.g. personality Briffa et al., 2015). Because of their reliance on individual assessment, rather than memory of past interactions, RHP-based hierarchies have been suggested to be particularly advantageous in open societies, where social interactions are drawn from a potentially large pool of individuals, thus limiting (de Silva et al., 2017b)—or alleviating (Boehm, 1999)—the ability to form hierarchies based on memory. For example, in Mandrills (*Mandrillus sphinx*) where groups can include up to 800 individuals, face colouration has been suggested to be a honest signal of dominance rank (Setchell et al., 2008), with the intensity of red colouration changing with changing status (Setchell & Dixson, 2001).

The second axis of research considers that the formation of dominance hierarchies requires individual recognition and memory of past interactions between individuals (Hobson, 2020). Dominance hierarchies based on memory are generally thought to only emerge in animals that form stable groups with relatively little turnover. This is because in more open societies, where social associations are drawn from a potentially large pool of individuals, the cognitive challenge of keeping track of many individuals over extended periods of time is thought to limit the ability for individuals to form hierarchies (de Silva et al., 2017a). In addition, if dominance hierarchies are also informed by third party interactions, the opportunity to observe those interactions may be more limited in open societies, further reducing the information available to individuals. Finally, open societies may alleviate the need for individuals to form hierarchies entirely, as individuals may be able to avoid repeated interactions by moving between groups (Boehm, 1999—but see Chaine et al., 2011, 2018; Penndorf et al., 2023).

Assumptions about limits to dominance hierarchies in large or open groups have recently been challenged by growing evidence that animals can form and maintain differentiated social relationships and complex multi-tiered social structures that can encompass tens or even hundreds of conspecifics (Papageorgiou et al., 2019; Papageorgiou & Farine, 2021; Camerlenghi et al., 2022). For example, in vulturine guineafowl (*Acryllium vulturinum*) that form multi-level societies (where groups interact with other groups), individuals maintain stable and steeply linear hierarchies in higher level social groupings that incorporate multiple breeding units (Dehnen, Papageorgiou, et al., 2022; Nyaguthii et al., 2025). In corvids that express fission-fusion dynamics, individuals have been shown to maintain long-term preferred social relationships that are likely based on individual recognition (Izawa & Watanabe, 2008; Chiarati et al., 2010; Boeckle & Bugnyar, 2012; Loretto et al., 2017; Boucherie et al., 2022). In this second case, it has been argued that such memory based interactions are represent a high degree of social complexity, and may have led to selection for social cognition (Bugnyar, 2013). However, the evidence remains limited to a relatively small number of well studied taxa, and more studies are needed to test (i) whether animals living in societies containing hundreds of potential social associates can form and maintain stable dominance hierarchies, and (ii) whether in species with high social mobility, individuals can be simultaneously part of several dominance hierarchies.

Parrots represent an excellent taxa in which to investigate these questions. Many parrot species exhibit strong pair bonds while also interacting within communal roosts and with individuals from the broader population. The outcome of this is that individuals may experience flock sizes ranging from several individuals to aggregations of thousands (Hardy, 1965; Noske et al., 1982; Rowley, 1990; O’Hara et al., 2019). Given this variable sociability, coupled with longevity (Wirthlin et al., 2018; Smeele et al., 2022) and complex cognition (Olkowicz et al., 2016), parrots are often referenced in discussions on social complexity (e.g., Krashenin-nikova et al., 2013; Hobson et al., 2014; Aplin et al., 2021). Yet studies of dominance patterns among wild parrots remain rare (but see Diamond and Bond, 1999; Penndorf et al., 2022), and it has often been assumed that parrot species which form large communal roosts cannot exhibit stable dominance hierarchies (Noske et al., 1982).

Studies of social interactions in parrots in captivity are more common, but results are mixed. Some species form linear or quasi-linear hierarchies (e.g., budgerigars, *Melopsittacus undulatus*: Soma and Hasegawa, 2004; cockatiels, *Nymphicus hollandicus*: Seibert and Crowell-Davis, 2001; Senegal parrot, *Poicephalus senegalus* : Lantermann, 1998) and others not (e.g., keas, *Nestor notabilis*: Tebbich et al., 1996, blue-fronted amazons, *Amazona aestiva* : de Souza Matos et al., 2017). The most detailed examination of this question comes from a series of captive studies on captive monk parakeets (*Myiopsitta monachus*), which revealed that groups of 11-21 individuals form linear social hierarchies based on memory of past interactions (Hobson et al., 2014; Hobson and DeDeo, 2015; Hobson et al., 2021; van der Marel et al., 2023). Furthermore, dominant individuals removed from their social group for a period of 8 days could not regain their rank immediately following reintroduction to the same group (van der Marel et al., 2023), providing further evidence that dominance ranks were the outcome of past interactions and memory, rather than the expression of intrinsic characteristics. However, because groups in captivity are necessarily much smaller and more stable than those in the wild, and captivity has been shown to induce the formation of linear hierarchies even in egalitarian species (Horová et al., 2015; but see Boucherie et al., 2022), questions remain about how these results translate to natural conditions. Wild studies are therefore vital to our understanding of dominance interactions in parrots and in open societies more generally.

Here, we quantify dominance hierarchies in and across a population of wild sulphur-crested cockatoos (SC-cockatoos; *Cacatua galerita*). SC-cockatoos are large, long-lived parrots that sleep in year-round communal roosts of 50-1000 birds that are the basis of social communities (Aplin et al., 2021). As roosts are located in areas that are also used for daily foraging, social interactions among individuals take place both at the roost (including during the day) and in the surrounding landscape as birds fission into small to medium foraging flocks (Noske et al., 1982; Styche, 2000; Aplin et al., 2021). Roost are open, and individuals can— and do—also regularly engage in between-roost movements (Aplin et al., 2021; Penndorf et al., 2022). Yet, despite these fission-fusion dynamics, SC-cockatoos maintain long-term social relationships beyond the pair bond that are strongly suggestive of social recognition, including with kin and non-relatives, and with birds from other roosts (Aplin et al., 2021; Penndorf et al., 2022).

In this study, we record social networks and aggressive interactions from 515 individuals across three neighbouring roosts to address three aims. First, we test whether social communities of wild SC-cockatoos that are centred on each roosting site form linear dominance hierarchies. Second, we identify individual predictors of dominance rank, including age and sex, and a proxy for resource holding potential, body weight. Third, we re-measure interactions across (i) two periods over two months (July and September 2019), and (ii) two periods over 3 years (2019-2022) to assess the short- and long-term stability of any emergent hierarchies.

## Methods

### Study population

The study was conducted at three neighbouring roost sites in north Sydney, Australia (Table S1). At each roost site, birds were habituated to the observer and then individually paint-marked with non-toxic dye (Marabu Fashion Spray, MARABU GmbH) using methods detailed in Penndorf et al. (2022). In addition to paint-marked birds, 144 birds were previously wing-tagged at the Royal Botanic Gardens as part of an ongoing citizen science study *Big City Birds* (Davis et al., 2017; Aplin et al., 2021). The number of marked birds at each roost varied from 42 - 165, depending on site and year (Table S1). Age (juveniles: <7 years, adults: >7 years) and sex of birds were assessed by eye-colour (Berry, 1981). Additionally, feathers were collected, from which DNA was extracted for molecular sexing and to match individuals across the two study periods.

Once marked, birds were trained to jump on a flat scale that read at 1g accuracy in exchange for a food reward (e.g. sunflower seed). This resulted in 214 birds being weighed across the three roost-sites in the first study period. Within individuals, weight was highly repeatable (0.78, 95% CI=0.72-0.82, R-package *rptR*, Stoffel et al., 2017, no. bootstraps=1000, *N*_*weightings per individual*_:1-17) and ranged from 717g to 1054g.

### Social data collection

Data on social associations and aggressive interactions were collected from all three sites in 2019 over two 10 day periods (July 8-20 and September 19 to October 2), resulting in a total of 165 observation hours. Data were then collected at one site (Clifton Gardens) in 2022 over 12 days (July 7-20) for a further 36 observation hours. During these periods, birds were attracted to forage on the ground by scattering small amounts of seed over an approximate 385-500*m*^2^ area of grass in parks close to the roost (300-680m distance). Foraging flocks were then observed daily for 2.5 to 3 hours. During each daily sample, the identity of all birds present—e.g. identifiable within the study area—was recorded every 10 minutes. Between presence scans, aggressive interactions were recorded using all occurrence sampling (Altmann, 1974). For each interaction, we recorded the time and the identities of winners and losers.

Presence scans were collated and used to construct social networks using a gambit-of-the-group approach and a simple ratio to weight edges between 0 (never observed in the same scan) and 1 (always observed in the same scan) (Cairns & Schwager, 1987; Farine & Whitehead, 2015; Hoppitt & Farine, 2018). We then identified social network communities using the fast-and-greedy algorithm included in the *igraph* package (Csardi, Nepusz, et al., 2006). These social network communities clearly mapped onto the roosting sites, and so we assigned the roost membership (residency) of each bird according to which social network community they were assigned to (Figure a).

### Dominance hierarchies

We calculated a separate dominance hierarchy for each of the three sites and observation periods. In order to obtain reliable dominance hierarchies, we only included individuals with seven or more agonistic interactions at a given roost-site (BA: N_*ind*_=144; CG: N_*ind*2019_=103, N_*ind*2022_=68; ; NB: N_*ind*_=82; Sánchez-Tójar et al., 2018—for more details about the choice of threshold used see Section S1.2, Figures S3, S4, S5). Hierarchies were calculated using randomized Elo-ratings (R-package *aniDom*, Sánchez-Tójar et al., 2018; sigma: 1/300, K: 200, randomisations: 10,000). We measured the robustness of the hierarchy by using the functions *‘repeatability by splitting’* and *‘repeatability by randomisation’*. Furthermore, we assessed the transitivity of the hierarchy following McDonald and Shizuka 2013 and Shizuka and McDonald 2015 (see Section S1.1).

To test whether dominance rank was predicted by age, sex, or body weight, we ran a Bayesian regression model using the R-package *brms* (beta-family, 4 chains, 4000 iterations, Bürkner, 2017a, 2017b, 2018). Since individuals could appear in the hierarchies of several sites (see Table S5) and across years, we included social community (BA, CG 2019, CG 2022, NB) and individual ID as random variables.

To assess the stability of dominance hierarchies over time, we calculated the hierarchy at each roost location separately for each observation period (July and September 2019 for short-term stability; 2019 and 2022 at Clifton Gardens for long-term stability). Using the *DynaRankR*-package (Strauss and Holekamp, 2019), we tested dyadic similarity between hierarchies between each time period at each site and each randomisation of the Elo-Rating (randomisation = 10,000). To calculate the random expectation, we created 10,000 hierarchies for each site and time period by randomly assigning dominance ranks to all individuals, before calculating the dyadic similarity obtained through random sampling. The resulting dyadic similarity score calculated by the *DynaRankR*-package takes into account the number of individuals within each hierarchy.

All analysis were conducted in R v 4.3.0 (R Core Team, 2023).

### Ethical Note

As the birds are never caught as part of our study, all participation is voluntary. While food provisioning may cause temporary changes in the cockatoos’ diet / behaviour, feeding is limited in time (only part of the day, for a maximum of 3 months a year), and is therefore not likely to induce a reliance on provisioning, or changes in their natural feeding behaviour given the longevity of the study species. Once the birds were habituated (hereafter defined as birds moving freely and without any signs of distress in close proximity to the observer), finger sponges (used to apply makeup) dipped in water-based, non-toxic, paint (MARABU Fashion Dye) were used to mark birds feeding within arms-reach. This method has been used previously (Klump et al., 2021; Penndorf et al., 2022), and has not been observed to elicit negative responses, as after marking: (i) birds stayed within close proximity of the observer, or—if startled by the touch/ movement— were back within seconds. (ii) birds did not change their behaviour (i.e. preening/ investigation of the paint-marks was not observed).

Once birds were marked, we opportunistically plucked 1-3 feathers per individual from the lower back, while individuals forage freely within arm-length of the observer. These feathers are used for genetic identification of individuals across years. This method has previously been employed in the species (Penndorf et al., 2022), and individuals (i) remained in proximity of the observer, and (ii) did not lose habituation as a result, suggesting that the impact was relatively low.

All procedures were approved by the ACEC (ACEC Project No. 19/2107), and were conducted under a NSW Scientific License (SL100107).

## Results

The number of birds included in the hierarchy at each site (i.e. ≥7 observed interactions) varied between 82 and 144 (Table S1). We recorded 6,402 (2019, 3 sites) and 2,087 (2022, 1 site) aggressive interactions across all birds, of which 5,694 and 2,006 respectively were between individuals that were included in the same dominance hierarchy (*N*_*ind* 2019_ = 202, *N*_*ind* 2022_ = 68). Most aggressive interactions between individuals of known sex were between males (*N*_*males* 2019_ = 72, *N*_*interactions males* 2019_ = 1,662, *N*_*interactions known sex* 2019_= 3,776, *N*_*males* 2022_ = 35, *N*_*interactions males* 2022_ = 633, *N*_*interactions knownsex* 2022_ = 1,479).

### Cockatoos form clear dominance hierarchies

We found that, within social communities, SC-cockatoos formed robust and highly transitive hierarchies (robustness: Figures 2a,b, S2 a,c,e,g & Table S2; transitivity ≥ 0.80—Table S4). Similar results were found without thresholding (i.e., including all individuals that interacted at least once at a given location—S1.5.1, robustness: Table S6; transitivity *geq*0.80, Table S7).

**Figure 1.**
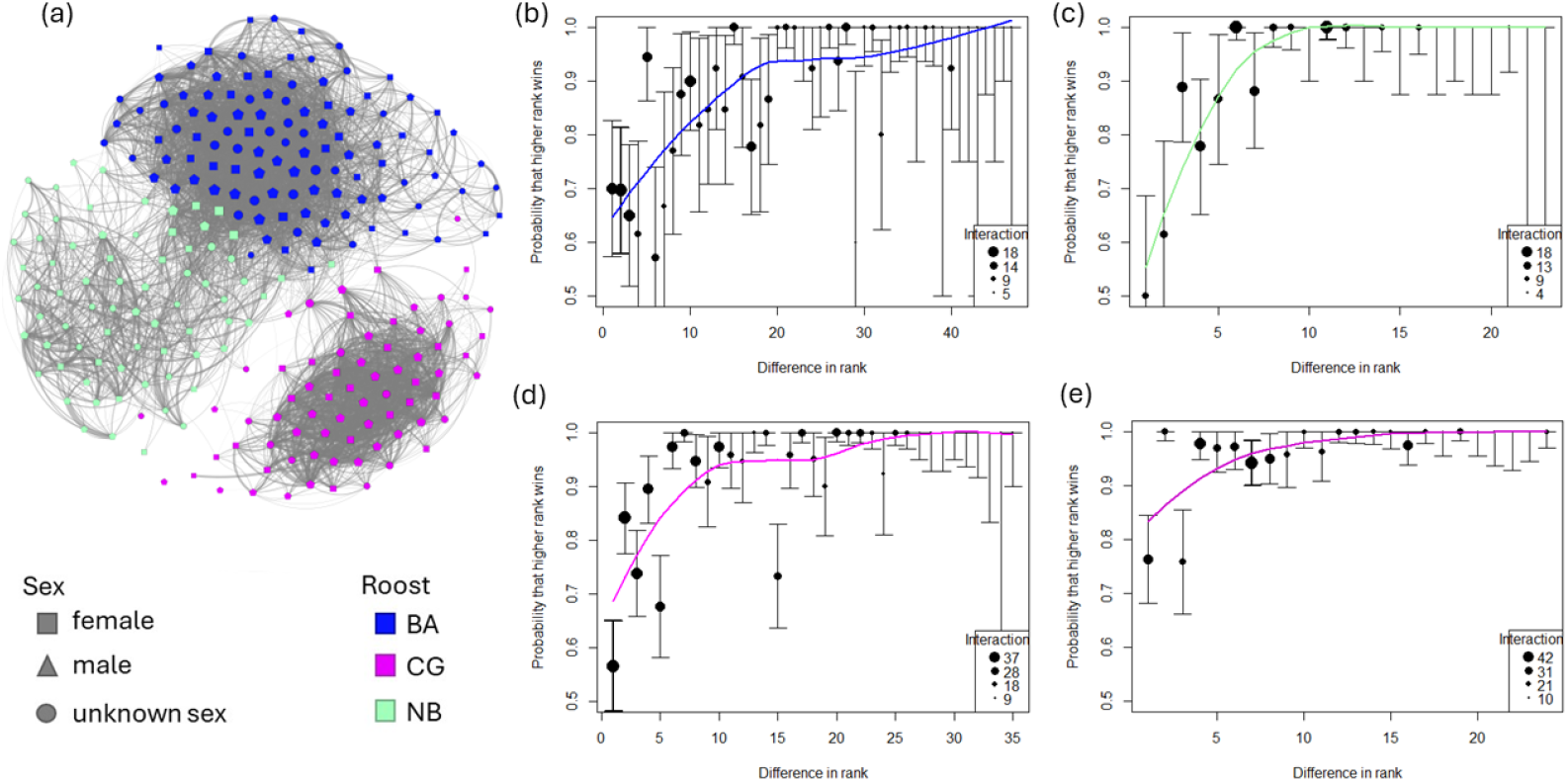
Social association network and steepness of dominance hierarchies. (a) Social network of sulphur-crested cockatoos in 2019. Node colour represents the three identified social communities identified using the “*fast-and-greedy* “ algorithm (blue: BA, pink: CG, green: NB). Node shape represents the sex of the individuals (square: female, triangle: males, circles: individuals of unknown sex). For visual clarity, only edges above 0.1 are represented. (b-e) The steepness of the male social hierarchy in three communities as identified in the social network (b: BA, c: NB, d-e: CG) across two years at CG (b-d: 2019, e: 2022). The error bars in b-d represent the standard deviation in elo-score for each individual across 10,000 randomisations.

**Figure 2.**
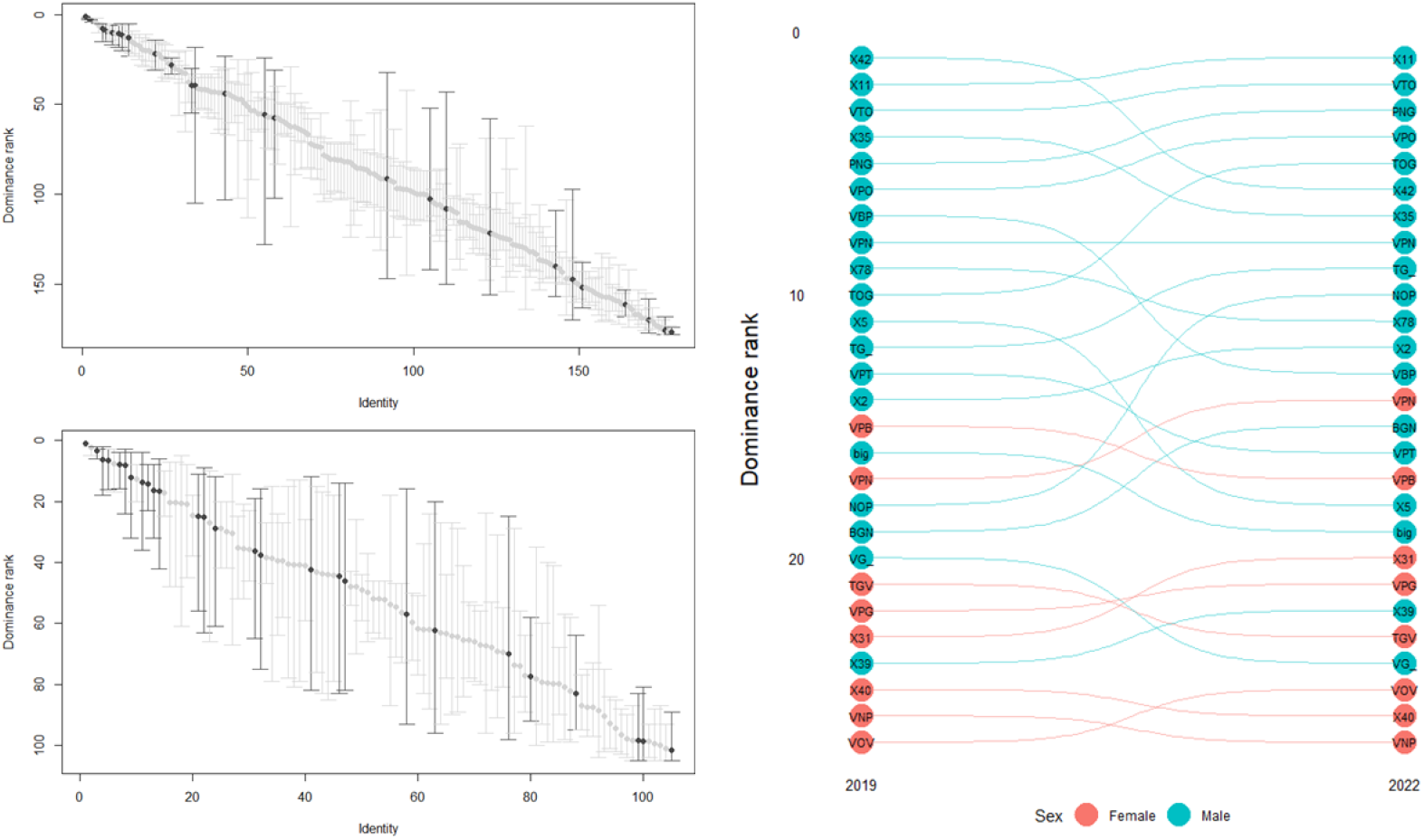
Mixed-sex dominance hierarchy at Clifton Gardens social community in (a) 2019 and (b) 2022. Individuals present in both study periods are shown in dark grey. Changes in their relative dominance rank across years is shown in (c). Males are represented in blue, and females in red.

### Predictors of dominance rank

Sex and age were significant predictors of rank, with males ranked higher than females (se:2.09, es: 0.24, 95%: [1.64, 2.57]—Figure S2 a,c,d,f), and adults ranked higher than juveniles (es: 0.82, se: 0.29, 95%CI: [0.26, 1.41]). However, body weight did not predict dominance rank (es: -0.61, se: 1.99, 95%CI: [-4.46, 3.32]). We obtain similar results when including all individuals that interacted at least once at a given site (i.e., no threshold, Section S1.5.2).

### Steepness of the hierarchy

Given the sex-based segregation of the dominance hierarchies, and that most interactions occurred within individuals of the same sex, we calculated the steepness of the hierarchies separately for each sex. In male hierarchies, individuals typically had a chance of >60% to win an aggressive interaction against a male one rank lower than themselves (Figure b-e). This probability increased relatively rapidly with increasing rank difference, reaching 0.9 with a rank difference between 4 and 16 (Figure b-e). In females (hierarchy only possible at Clifton Gardens in 2019 and 2022), the probability of winning against an individual of one rank below or above was more variable (2019: 0.6, 2022: 0.85), and reached 0.9 for a rank difference of 8 and 3 respectively (Figure S1a,b—for values of steepness without threshold, see S1.5.3).

### Stability of dominance hierarchies over time

Within each social community, dominance ranks were highly repeatable over a period of two months (July-September 2019; dyadic similarity >0.77, Table S4), which was significantly higher than expected by chance (R ≈ 0.5, Table S4). Similar repeatability was found when repeating the analysis within each age (juvenile/adult) and sex (males/females) class (Table S4), suggesting that this effect was not driven by predictable age and sex differences in dominance rank. In the CG social community measured again after a period of 3 years, dominance ranks were very similar in their repeatability (2019-2022, similarity: 0.84 [0.81-0.87], random expectation: 0.50 [0.42-0.59], N=27, Figure 2c, Table S4). Similar values of dyadic similarity were found when not using any threshold (i.e., including all individuals that interacted at least once—Section S1.5.4 & Table S8).

## Discussion

We found that, despite their largely open social structure and social communities of potentially hundred of individuals, SC-cockatoos formed clear linear dominance hierarchies. These hierarchies were structured by sex and age, with males and adults tending to be higher in the hierarchy than females and juveniles. However, while males tend to be larger than females, there was no evidence for an effect on body size on rank. Finally, SC-cockatoos were consistent in dominance across time: hierarchies were highly stable across periods of months to years.

### RHP and the determination of dominance ranks in SC-cockatoos

Body weight—a measure of body size that is often linked to resource-holding potential (reviewed by Arnott and Elwood, 2009)—did not predict dominance ranks. Yet, SC-cockatoos maintain stable dominance hierarchies across months and years. This leads to several speculations about the mechanisms of hierarchy formation and maintenance in this species (reviewed by Dehnen, Arbon, et al., 2022; Tibbetts et al., 2022). First, hierarchies in this system could still be based on RHP. Body weight constitutes only one of the potential indicators of resource holding potential (Allen & Krofel, 2022), and we cannot exclude that there is another, undescribed status signal in this species. For example, colouration (Santos et al., 2011; Beltrao et al., 2021) is an indicator of dominance rank in many species, while crest-length (Dakin, 2011) is an indicator of dominance in Indian peafowl (*Pavo cristatus*). SC-cockatoos are almost entirely white except for their large yellow crests, which are used in signalling, for example at nest hollows. However, we believe it to be unlikely that crest-colouration or length signal dominance in our study, as crest-displays were only performed in 1 % of the dominance interactions that we recorded.

Second, in the absence of physical predictors of dominance ranks, it is possible that aggressive interactions in this species are primarily based on social recognition and memory of past interactions. If the formation and maintenance of dominance hierarchies in this species requires individuals to recognize and remember all members of their social communities, SC-cockatoos in our population may need to remember at least 68 individuals with high daily turnover, local hierarchies comprised between 68 and 144 individuals. This would constitute a memory for conspecifics that is comparable to that of ravens (Boucherie et al., 2022), elephants (McComb et al., 2000) and humans (Hill and Dunbar, 2003, but see Lindenfors et al., 2021). This seems possible, even plausible, given that SC-cockatoos also share other life-history traits with these species, including large relative brain-size and extended longevity (Smeele et al., 2022), and that this would be in line with captive studies in monk parakeets (Hobson et al., 2014; Hobson & DeDeo, 2015; Hobson et al., 2021). Further exploration will be needed to fully test the potential for reliance on memory of past interaction in this system.

The third possible mechanism to explain the existence of orderly hierarchies in such large groups is that individual recognition and memory of a small subset of individuals (e.g., individuals of the same sex, or frequent associates, see Chaine et al., 2011, 2018) is sufficient to maintain a stable, linear, dominance hierarchy at the scale of social communities. We found that SC-cockatoos exhibit a sex- and age-segregated hierarchy, with males above females and adults above juveniles. In this case, a categorisation of dominance relationships (i.e. only remembering the exact dominance ranks of conspecifics of the same age class and sex) would significantly reduce cognitive requirements that come with making social decisions in an open society with significant fission-fusion dynamics (Aureli & Schino, 2019). In our study, this heuristic would reduce the requirement for individual recognition substantially; for instance, individual adult males in our population could encounter between 19 (1 site, NB) and 72 (all sites) other adult males on any given day.

It should also be noted that our roost sizes, while typical for our study population, are relatively small for SC-cockatoos. Roost sizes in this species have been reported as reaching well over one thousand individuals (Aplin et al., 2021), and in theses cases, even a sex and age based categorisation would still result in very large numbers of potential interaction partners. Further study is needed to determine what strategies SC-cockatoos might use to reduce the cognitive demands of living in such enormous and dynamic societies, and whether stability and/or linearity of hierarchies break down at larger social scales.

### Stability in rank over space and time

Social communities were clearly differentiable in our social network, yet social communities were also connected, with a subset of individuals exhibiting high social mobility (Penndorf et al., 2022). As a result, some individuals in our study were present in two or three dominance hierarchies, and were likely to be familiar with individuals across a much broader social landscape than just their own social community. Dominance hierarchies in the three social communities measured were temporally stable when re-measured after a period of two months. The dominance hierarchy was further re-measured in one social community after a period three years, and showed a similarly level of repeatability. Together, this suggests that dominance hierarchies are highly stable over at least three years, and perhaps even over longer periods of time

This social stability is consistent with some previous research—although higher than previously found in birds. Common ravens, for example, form communal roosts with high fission-fusion dynamics similar to SC-cockatoos, yet the social stability in the social group (3 % across 7 years—Boucherie et al., 2022, measured as percentage of individuals present in both time periods) is around sixteen times lower than the one observed in our study (30% across 3 years). However, stable dominance hierarchies have been shown in other open systems with fission-fusion dynamics. For example, female mountain goats (*Oreamnos americanus*) form linear hierarchies based on age, that are highly stable from one year to the next (*R*^2^ *>*0.93) (Côté, 2000), and similar measures of stability in dominance ranks have been suggested for other ungulate species (Hirotani, 1990). Taken together, the social stability observed in SC-cockatoos is therefore remarkable, though not unheard of in species with high fission-fusion dynamics. This is perhaps unsurprising; SC-cockatoos are long-lived birds with an extended juvenile period of 7-years and adult lifespans of several decades (Smeele et al., 2022). Previous research on SC-cockatoos found that birds tend to maintain differentiated social relationships over periods of at least 18 months, with these relationships becoming more stable with age as measured over 10 years (Aplin et al., 2021). Our study did not allow for an similar analysis of age, but future research could ask how such increasing social stability influences dominance rank and relationships.

## Conclusion

Wild SC-cockatoos can form stable long-term relationships (Aplin et al., 2021), and maintain some social relationships even after movement into different social communities (Penndorf et al., 2022). Our results build upon these findings by indicating that SC-cockatoos also form and maintain clear and stable dominance relationships with a large number of conspecifics, and retain their dominance rank over periods of at least three years. Taken together, these studies provide ample evidence these long-lived parrots exhibit hidden social stability within these extended and dynamic social networks. It adds to evidence from other large-brained bird species that exhibit fission-fusion dynamics, such as common ravens (Boeckle and Bugnyar, 2012), that such species possess social cognitive abilities that allow them to navigate a extended social landscape with within- and between-community social interactions. Future studies should attempt to elucidate the cognitive load of maintaining dominance hierarchies in such systems as well as the limits of such social structures, contributing to ongoing debates about the evolution and ecology of socio-cognitive complexity.

## Acknowledgements

We acknowledge the Gamaragal and Gadigal people as the Traditional Custodians of the Land on which this study was conducted. We thank two anonymous reviewers for their insightful comments. All procedures were approved by the ACEC (ACEC Project No. 19/2107), and were conducted under a NSW Scientific License (SL100107). This work was supported by the Swiss State Secretariat for Education, Research and Innovation (SERI) under contract number MB22.00056. Part of the work was supported by funding from the Max Planck Society to JP and LMA. DRF was funded by an Eccellenza Professorship Grant of the Swiss National Science Foundation (Grant Number PCEFP3 187058) and the European Research Council (ERC) under the European Union’s Horizon 2020 research and innovation programme (grant agreement No. 850859).

