## Supplementary material for "Long-term stability of dominance hierarchies in a wild parrot with fission-fusion dynamics"

#### Electronic Supplementary Materials

### S1 Methods

#### S1.1 Transitivity of the hierarchy

In order to assess the transitivity of the hierarchy, we used a directed 1/0 matrix, based on the majority of wins (McDonald and Shizuka, 2013), with the modification that when two individuals won the same number of contests against each other, they both received scores of 1 (Shizuka and McDonald, 2015). This produced a directed network where a dominant-subordinate relation was represented by an edge directed from the dominant to the subordinate individual. We used the package *compete* (Curley, 2016) to infer transitivity of dominance relationships both at group level, and within each sex. Emphasis was on the proportion of transitive triangles relative to all triangles (Pt, defined by McDonald and Shizuka, 2013) is given by:

$$P_t = \frac{N_{transitive}}{N_{transitive}+N_{cycle}}$$

where $N_{transitive}$ is the number of transitive dyads, and $N_{cycle}$ is the number of cyclic triangles. In a random network, $P_t$ is expected to be 0.75 (McDonald and Shizuka, 2013). Therefore, transitivity was scaled so that it runs from 0 for the random expectation to 1 (all triangles are transitive), by the following triangle transitivity metric ($t_{tri}$):

$$t_{tri} = 4 \times (P_t - 0.75)$$

#### S1.2 Choice of threshold

Sánchez-Tójar et al. (2018) suggested that a ratio of observed interactions to individuals of at least 10 is needed for a reliable inference of dominance hierarchies. Therefore, we calculated the number of interactions and individuals in our dataset with varying the threshold (minimum number of interactions for an individual to be included). We found that the shape of the mean ratio of interactions per individual is non-monotonic. It increases sharply as we threshold individuals with few observations and the increase slows as we threshold individuals with more observations (Figure S4). Thus, increasing the threshold is initially beneficial, but these benefits decrease as the threshold value gets larger. A threshold of 10 appears to match the point where the increase slows.

In a second step, we calculated the dyadic similarity (Strauss & Holekamp, 2019) of hierarchies calculated with thresholds varying from 1 and 9, and the hierarchy calculated with a threshold of 10 (Figure S6). We found that the shape of the increase in the correlation between the inferred hierarchy for a given threshold and for the threshold of 10 interactions per individual is monotonic, meaning that there is no threshold that is necessarily better than another. Finally, we found a sharp decrease in the number of individuals included in the hierarchy as a function of the threshold.

Together, these suggest that a threshold of 7 interactions per individual represents a good balance between robustness of the hierarchy, maximising the ratio of interactions to individuals, and including sufficient individuals in our analyses.

#### S1.3 Do we have sufficient data?

To determine whether the number of interactions collected at each site (BA, CG, and NB) and each year (CG 2019, CG 2022) was sufficient to infer reliable dominance hierarchies, we subsetted the dataset by randomly sampling 10, 20, 30, 40, 50, 60, 70, 80 or 90% of all interactions. For each subset, we inferred the dominance hierarchy, including only individuals with at least 7 aggressive interactions at the site (for details on the choice of threshold, see S1.2). We then compared the dominance ranks inferred from the subsetted dataset to the dominance ranks inferred from the full dataset, by calculating the dyadic similarity (package *DynaRankR*, Strauss and Holekamp, 2019). We repeated the procedure 100 times for each subset.

We find that the dyadic similarity at each site stabilizes when using 70 (CG 2019 2022) to 80% (BA, NB) of our data, suggesting that our dataset is sufficient to infer reliable dominance hierarchies (Figure S11).

#### S1.4 Robustness of the dyadic similarity

To check whether our dataset allowed a robust estimation of the similarity of social hierarchies between periods (July-September 2019—3 sites) or years (2019-2022—1 site), we :

1. randomly selected 30, 40, 50, 60, 70, 80 or 90 % of all interactions recorded at each site,
2. calculated the local hierarchies for each period (July-September 2019, or 2019-2022, see Section "Dominance hierarchies" in the main manuscript), without thresholding—i.e., including all individuals that interacted at least once at any given site,

3. calculated the dyadic similarity between the two periods, as well as the random dyadic similarity (Strauss, 2020).

4. We then repeat steps one to three 200 times.

We found that dyadic similarities between periods calculated with as as little as 30% of our dataset significantly above random (Figure S7).

#### S1.5 Impact of threshold on results

We rerun all analysis presented in the main manuscript, but without thresholding based on the number of interactions, i.e., including all individuals recorded interacting at least once. All methods are described in the Methods section of the main manuscript.

##### S1.5.1 Cockatoos form repeatable and transitive dominance hierarchies

We found that, within social communities, SC-cockatoos formed robust and highly transitive hierarchies (robustness: $> 0.67$, Table S6; transitivity $\geq 0.65$—Table S7).

##### S1.5.2 Predictors of dominance rank

We find again that sex and age were significant predictors of rank, with males ranked higher than females (se:1.84, es: 0.24, 95%: [1.38,2.32]), and adults ranked higher than juveniles (es: 0.68, se: 0.27, 95%CI: [ 0.16, 1.23]). However, body weight did not predict dominance rank (es: -0.17, se: 1.99, 95%CI: [-4.12, 3.72]).

##### S1.5.3 Steepness

In male hierarchies, individuals typically had a chance of >60% to win an aggressive interaction against a male one rank lower than themselves (Figure S9b-e). However, this probability increased relatively rapidly with increasing rank difference, reaching 0.9 with a rank difference between 4 and 8 (Figure S9). In females (hierarchy only possible at Clifton Gardens in 2019 and 2022), the probability of winning against an individual of one rank below or above was more variable (2019: 0.6, 2022: 0.65), and reached 0.9 for a rank difference of 12 (2022) or 15 (2019) (Figure S10a,b).

##### S1.5.4 Stability of dominance hierarchies over time

Within each social community, dominance ranks were highly repeatable over a period of two months (July-September 2019; Table S8), which was significantlyhigher than expected by chance (R $\approx$ 0.5, Table S8). Similar repeatability was found when repeating the analysis within each age (juvenile/adult) and sex (males/females) class (Table S8), with the exeption the hierarchies of females and juveniles at one site (Table S8). In the CG social community measured again after a period of 3 years, dominance ranks were very similar in their repeatability (2019-2022, Table S8).

#### S2 Figures

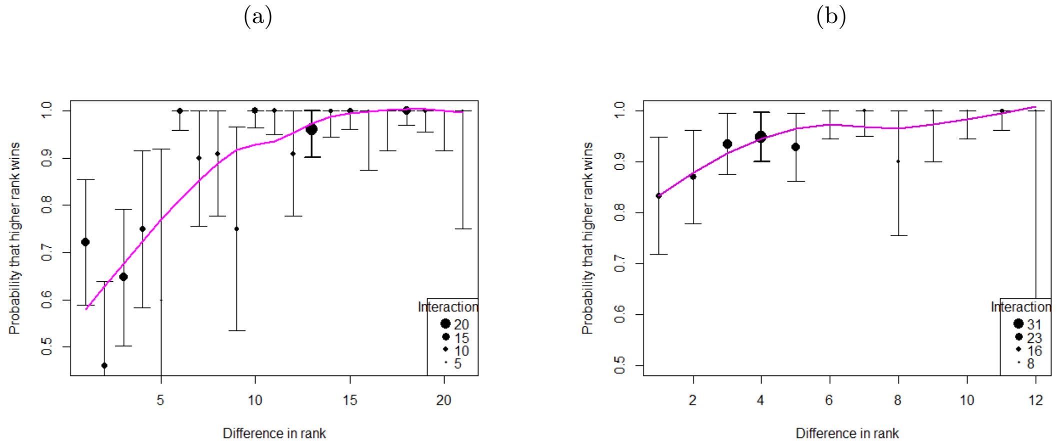


Figure S1: Shapes of female hierarchies measured at the CG-roost across two years: a: 2019, b: 2022. Only individuals with at least 7 interactions at a given site were included.

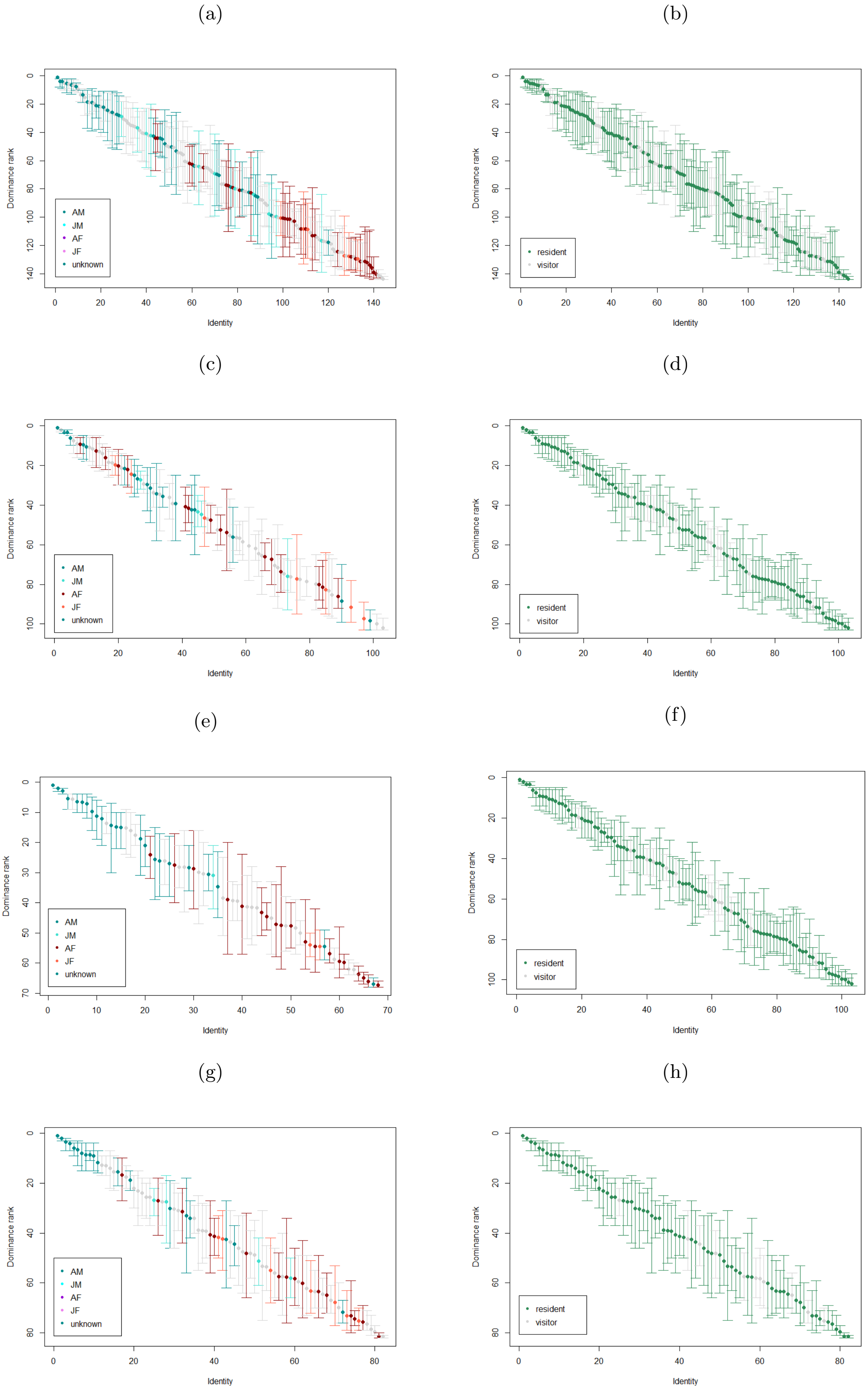


Figure S2: Hierarchies coloured by age and sex (left column) or residence status (right column) in three roosting communities of SC-cockatoos: a-b: Balmoral Beach (BA), c-d: Clifton Gardens (CG) 2019, e-f: Clifton Gardens 2022, g-h: Northbridge (NB).
AM: adult male, JM: juvenile male, AF: adult female, JF: juvenile female. All individuals for whom information about age and sex were incomplete or absent are represented in grey (left column).

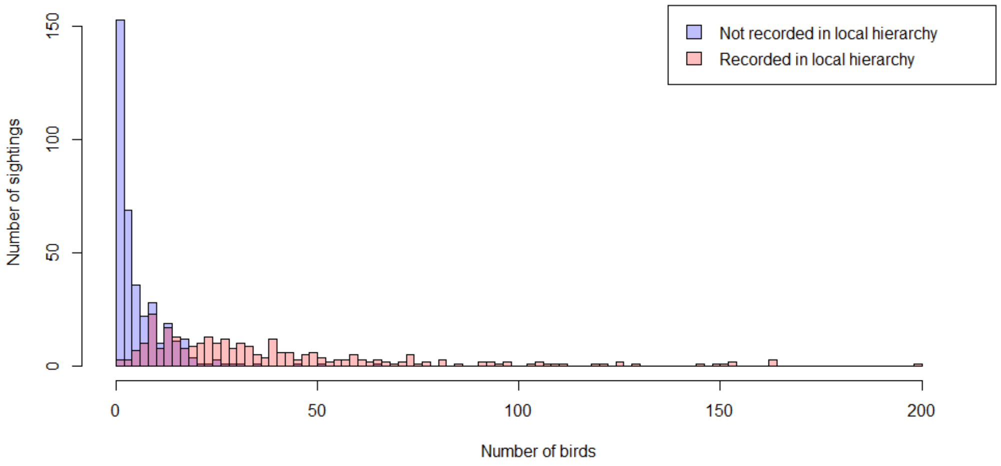


Figure S3: Histogram representing the number of sightings at one of the study sites as function of the number of birds sighted. The blue bars represent birds that were sighted at a site, but which were not include in the local hierarchy (less than 10 interactions at the site). The red bars represent individuals that were site at a site, and included in the local hierarchies (more than 10 interactions at the site). Individuals visiting several times are represented once at each site they visited.

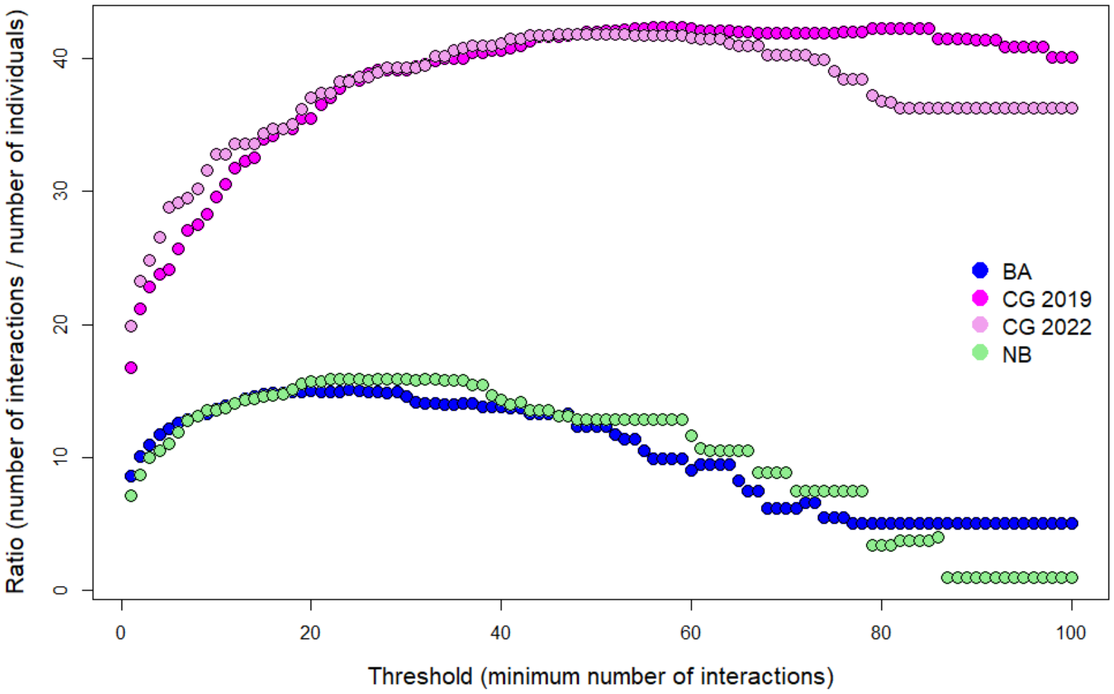


Figure S4: Ratio number of interactions in the dataset per number of individuals as function of the threshold(the minimum number of interactions for individuals to be included).

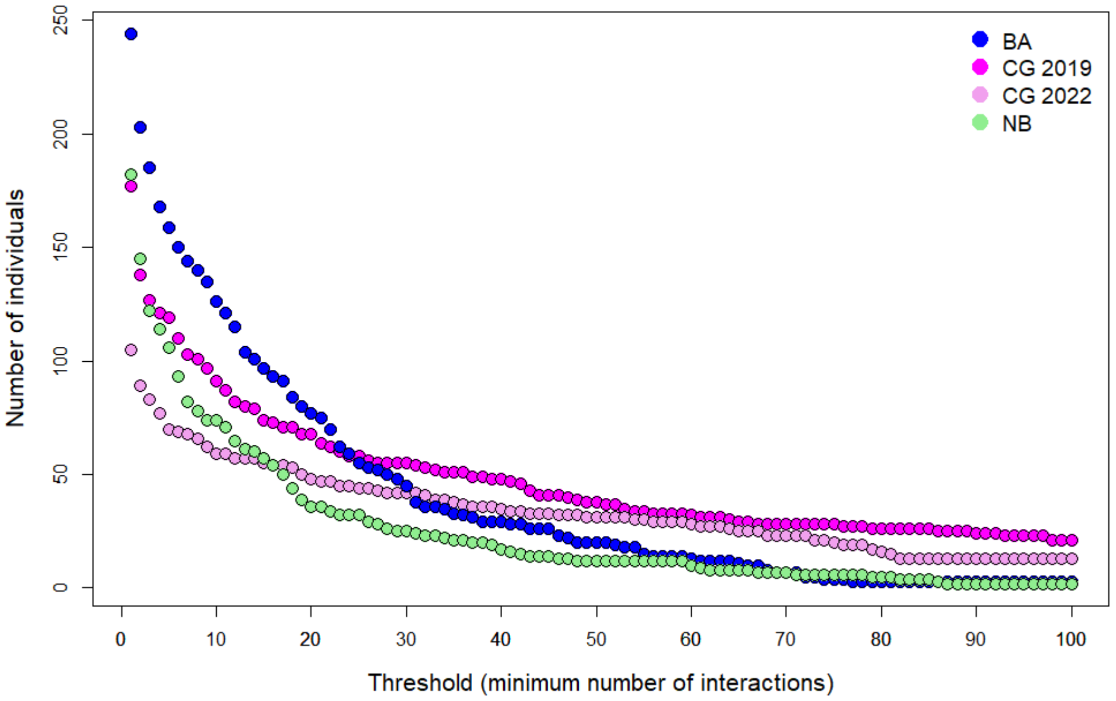


Figure S5: Number of individuals included in the hierarchy at each site, depending on the threshold (minimum number of interactions at each site).

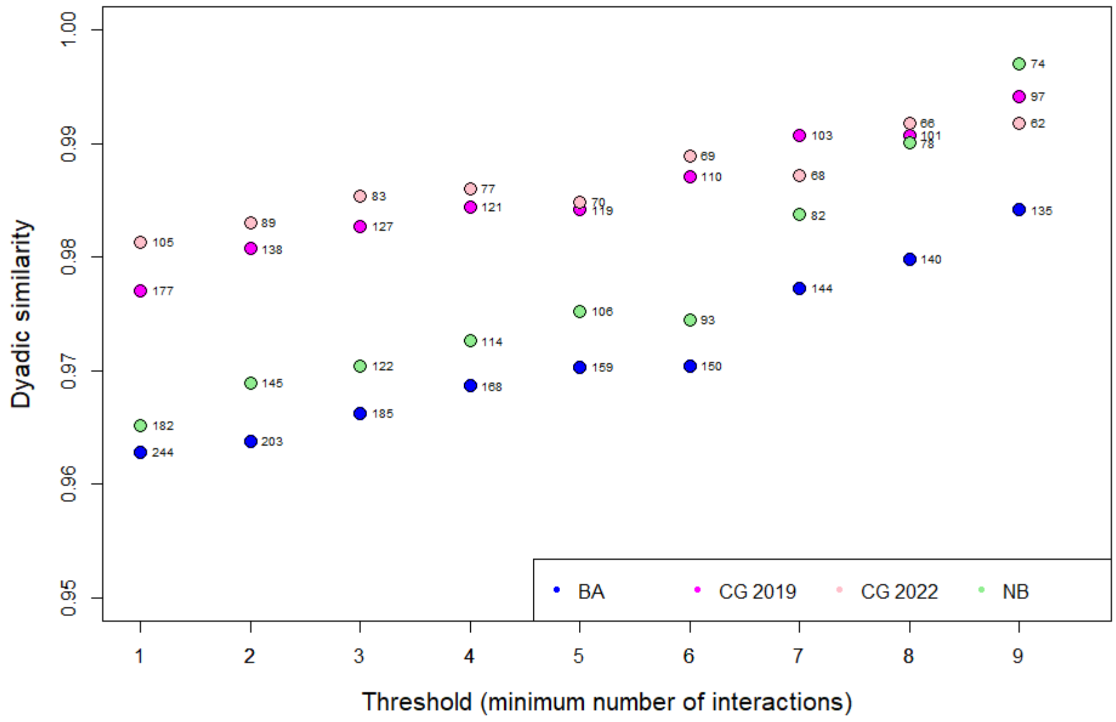


Figure S6: Dyadic similarity calculated between hierarchies calculated with varying thresholds (x-axis) and a hierarchy calculated including only interactions with at least 10 interactions. The numbers represent the number of individuals included in each hierarchy.

(a) (b)

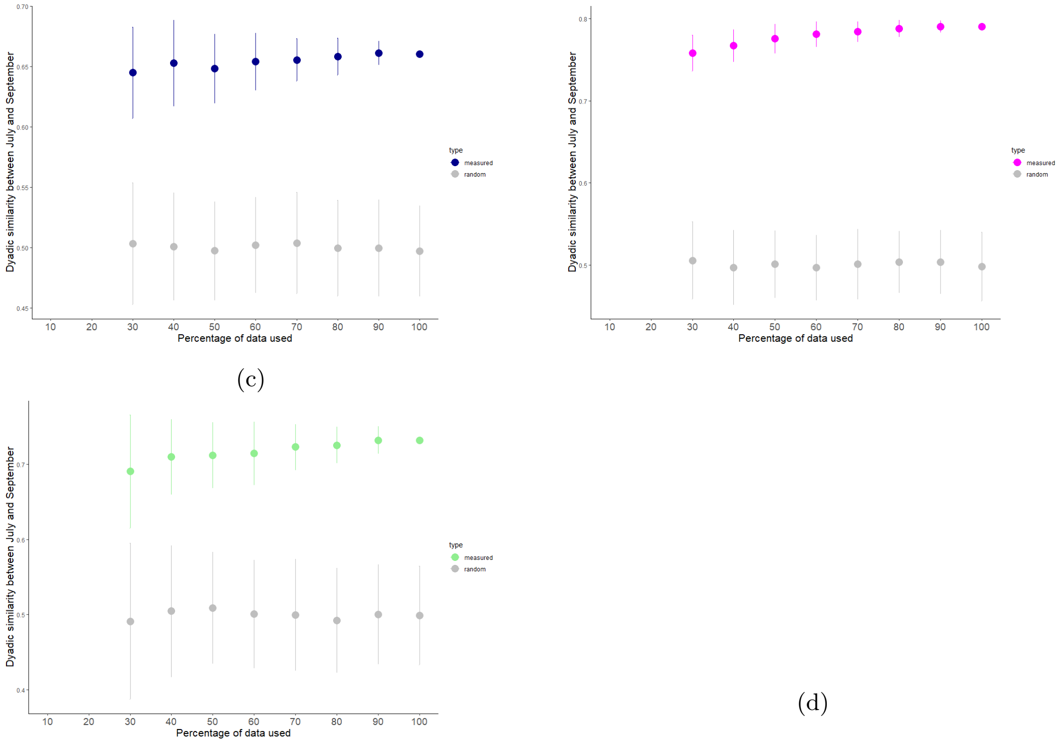


(c) (d)

Figure S7: Dyadic similarity between hierarchies in July and September, as function of the percentage of the data used to calculate the hierarchies, calculated at each site: (a) Balmoral, (b) Clifton Garden, and (c) Northbridge. The circles represent the mean dyadic similarity and the error bars the standard deviation, calculated across 200 randomisations.

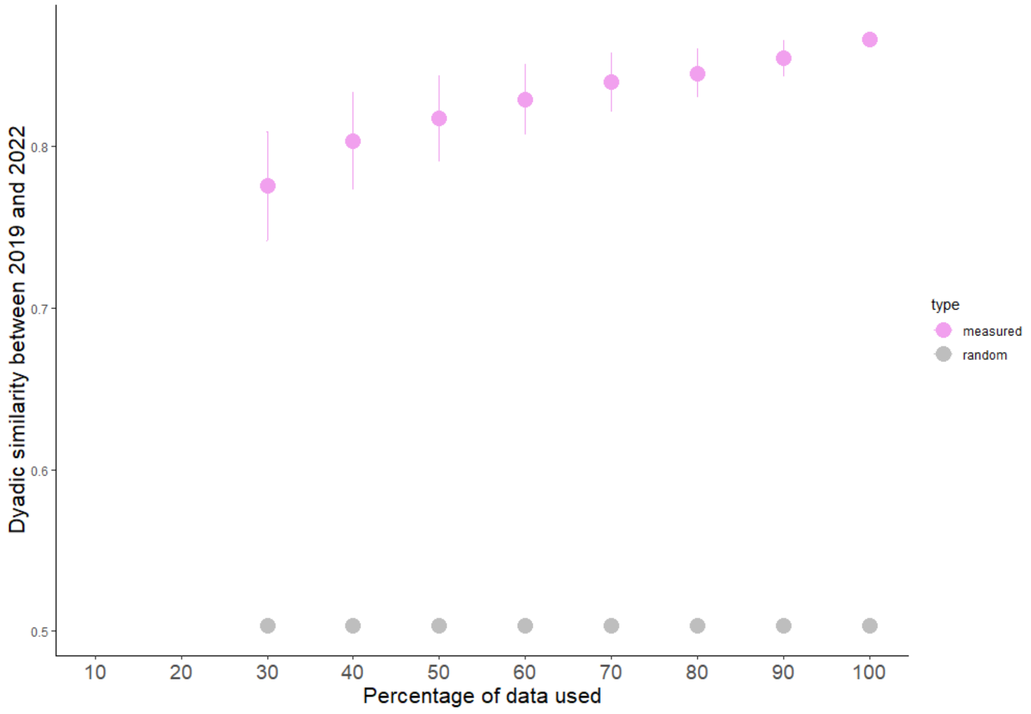


Figure S8: Dyadic similarity between hierarchies recorded at CG in 2019 and 2022, as function of the percentage of the data used to calculate the hierarchies. The circles represent the mean dyadic similarity and the error bars the standard deviation, calculated across 200 randomisations.

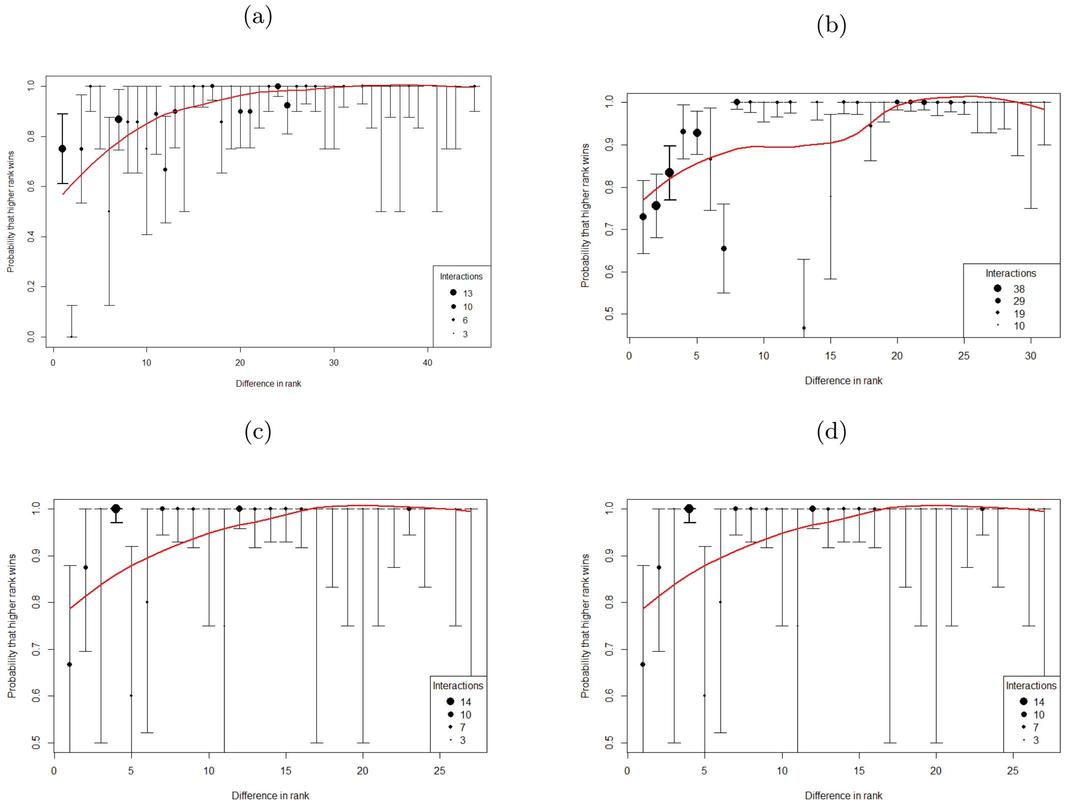


Figure S9: Shapes of male hierarchies, including all males observed interacting at least once at one of our study sites: (a) BA, (b) CG 2019, (c) CG 2022, (d) NB

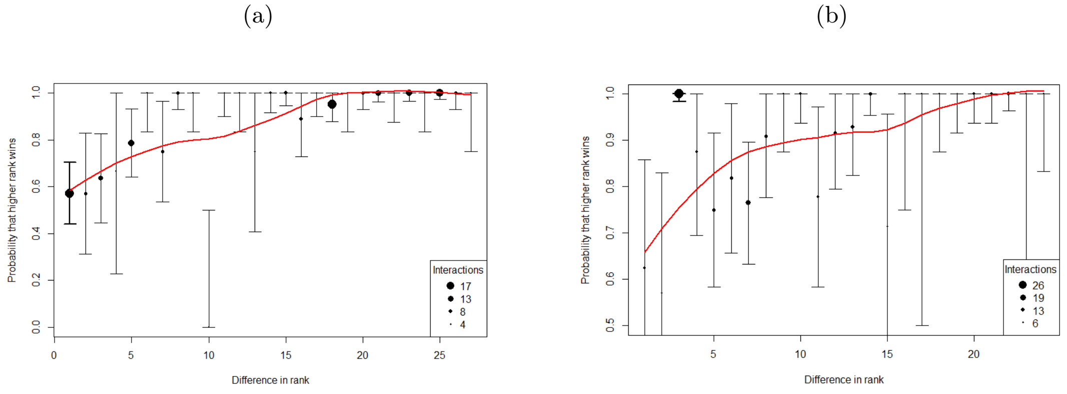


Figure S10: Shapes of female hierarchies, including all females observed interacting at least once at CG in 2019 (a) or 2022 (b).

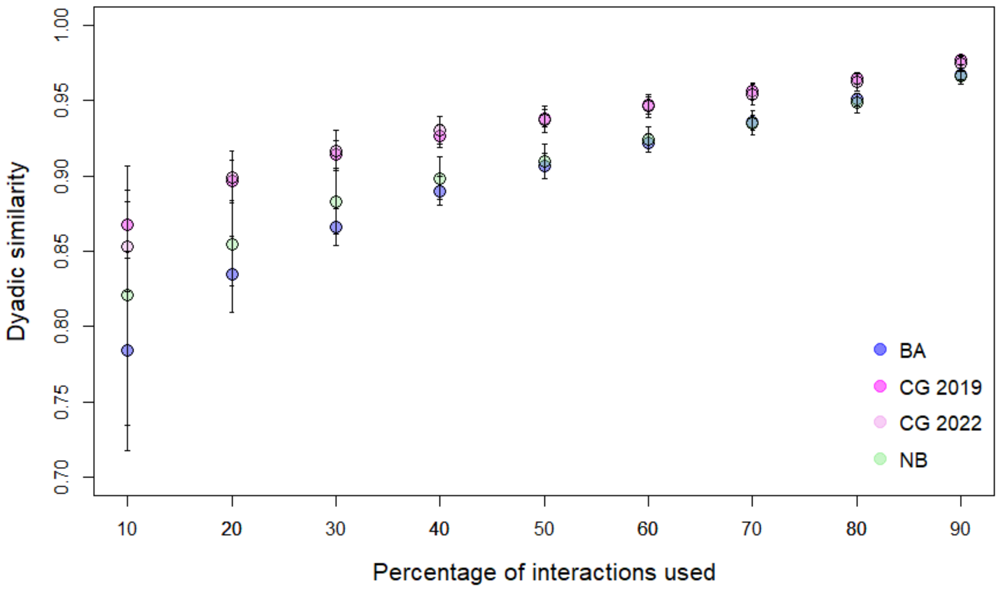


Figure S11: Dyadic similarity between dominance hierarchies inferred on a subset of the dataset (10 - 90%) of the data, and dominance hierarchies inferred on the full dataset. Points represent the mean similarity score, and bars the standard error.
All hierarchies only include individuals with at least 7 interactions in the subsetted dataset.

#### S3 Tables

Table S1: Location of the three study roosts and the estimated number of SC-Cockatoos at each site. The two letter code between parentheses represents the abbreviation for each roost site. At each site, we conducted roost counts by attracting birds to the ground at dawn, and recording the number and identities of all birds present. These roost counts were then used to estimated the number of birds at each site, as well as the percentage of marked birds (2019). As individuals regularly visit other different sites, the number of birds marked at one specific site can be larger than the number of individuals roosting at this location. P: paintmarked; W: wingtagged; R: recognizable feature. For Northbridge, the number between brackets represent the number and percentage of marked individuals of the satellite roost.

| Year | Roosting site (abbreviation) | GPS location | Number of marked birds | Roost size | Percentage marked |
|---|---|---|---|---|---|
| 2019 | Balmoral Beach (BA) | -33.828494, 151.253983 | 209P, 12W, 6R | 88-165 | 94-96% |
| 2019 | Clifton Garden (CG) | -33.841444, 151.252889 | 86P, 12W, 1R | 42-111 | 92-95% |
| 2022 | Clifton Garden (CG) | -33.841444, 151.252889 | 92P, 13W, 0R | NA | NA |
| 2019 | Northbridge (NB) | -33.817167, 151.221694 | 78P, 2W, 4R | 56-72 (+21-27) | 93-98 % (100%) |

Table S2: Uncertainty of SC-cockatoo social hierarchies measured at each of the three roost sites. Correlations above 0.8 (repeatability by randomisation) and 0.5 (repeatability by splitting) suggest robust hierarchies (Sánchez-Tójar et al., 2018)

| Year | Roosting site | Repeatability by randomisation | Repeatability by splitting [CI] | $N_{individuals}$ | $N_{interactions}$ |
|---|---|---|---|---|---|
| 2019 | BA | 0.96 | 0.78 [0.73, 0.83] | 144 | 1.856 |
| 2019 | CG | 0.97 | 0.86 [0.82, 0.90] | 103 | 2.790 |
| 2022 | CG | 0.97 | 0.83 [0.77, 0.89] | 68 | 2.006 |
| 2019 | NB | 0.97 | 0.74 [0.66, 0.81] | 82 | 1.048 |

Table S3: Transitivity of aggression networks at SC-cockatoo group level, calculated for all individuals present at one site (overall), or within each sex.

| Year | Roosting site | Overall | Males | Females |
|---|---|---|---|---|
| 2019 | BA | 0.80 | 0.69 | / |
| 2019 | CG | 0.89 | 0.99 | 0.73 |
| 2022 | CG | 0.88 | 0.99 | 0.74 |
| 2019 | NB | 0.87 | 0.94 | / |

Table S4: Dyadic similarity in social hierarchy within months (July and September 2019) or years (2019-2022), calculated using the *DynaRankR*-package (Strauss & Holekamp, 2019)

Dominance ranks were calculated using randomized elo-ratings, and only individuals with at least 7 interactions at a site were included in the analysis.

| Level | Time period | Roosting site | Similarity | Random expectation |
|---|---|---|---|---|
| All ages and sexes | July & September 2019 | BA | 0.77 [0.75, 0.80] | 0.50 [0.42, 0.58] |
| | July & September 2019 | CG | 0.83 [0.82, 0.84] | 0.50 [0.45, 0.55] |
| | July & September 2019 | NB | 0.98 [0.91, 1] | 0.50 [0.18, 0.82] |
| | 2019 & 2022 | CG | 0.84 [0.81, 0.87] | 0.50 [0.42, 0.59] |
| Juveniles | July & September 2019 | BA | / | / |
| | July & September 2019 | CG | 0.81 [0.76, 0.85] | 0.50 [0.36, 0.65] |
| | July & September 2019 | NB | / | / |
| | 2019 & 2022 | CG | / | / |
| Adults | July & September 2019 | BA | 0.85 [0.82, 0.89] | 0.50 [0.37, 0.64] |
| | July & September 2019 | CG | 0.86 [0.85,0.88] | 0.50 [0.43, 0.57] |
| | July & September 2019 | NB | / | / |
| | 2019 & 2022 | CG | 0.86 [0.83, 0.89] | 0.50 [0.41, 0.59] |
| Females | July & September 2019 | BA | / | / |
| | July & September 2019 | CG | 0.66 [0.60, 0.71] | 0.50 [0.40, 0.60] |
| | July & September 2019 | NB | / | / |
| | 2019 & 2022 | CG | 0.85 [0.78, 0.93] | 0.50 [0.32, 0.67] |
| Males | July & September 2019 | BA | 0.82 [0.77, 0.87] | 0.50 [0.36, 0.63] |
| | July & September 2019 | CG | 0.84 [0.82, 0.87] | 0.50 [0.42, 0.58] |
| | July & September 2019 | NB | / | / |
| | 2019 & 2022 | CG | 0.80 [0.74, 0.84] | 0.50 [0.38, 0.62] |

Table S5: Table summarizing the number of roosts visited by individuals, and the number of local hierarchy they are part of. The number of visited roosts is based on group scans at the 3 different in which the individuals appeared, and the inclusion within the local hierarchies on whether the individuals were involved at least 7) aggressive interactions at this site.

| Number of roosts visited | Number of hierarchies | Number of individuals |
|---|---|---|
| 3 | 0 hierarchies | 5 |
| | 1 hierarchy | 14 |
| | 2 hierarchies | 12 |
| | 3 hierarchies | 2 |
| 2 | 0 hierarchies | 36 |
| | 1 hierarchy | 78 |
| | 2 hierarchies | 25 |
| 1 | 0 hierarchies | 133 |
| | 1 hierarchy | 154 |

Table S6: Uncertainty of SC-cockatoo social hierarchies measured at each of the three roost sites, including all individuals observed interacting at least once. Correlations above 0.8 (repeatability by randomisation) and 0.5 (repeatability by splitting) suggest robust hierarchies (Sánchez-Tójar et al., 2018).

| Year | Roosting site | Repeatability by randomisation | Repeatability by splitting (CI) | $N_{individuals}$ | $N_{interactions}$ |
|---|---|---|---|---|---|
| 2019 | BA | 0.96 | 0.74 (0.68 - 0.79) | 244 | 2,090 |
| 2019 | CG | 0.97 | 0.82 (0.77 - 0.86) | 187 | 2,959 |
| 2022 | CG | 0.97 | 0.80 (0.73 - 0.86) | 105 | 2,087 |
| 2019 | NB | 0.97 | 0.67 (0.60 - 0.75) | 182 | 1,299 |

Table S7: Transitivity of aggression networks at SC-cockatoo group level, calculated for all individuals present at one site (overall), or within each sex. All individuals observed interacting at least once where included (no threshold).

| Year | Roosting site | Overall | Males | Females |
|---|---|---|---|---|
| 2019 | BA | 0.80 | 0.69 | / |
| 2019 | CG | 0.89 | 0.85 | 0.65 |
| 2022 | CG | 0.88 | 0.99 | 0.74 |
| 2019 | NB | 0.88 | 0.93 | / |

Table S8: Dyadic similarity in social hierarchy within months (July and September 2019) or years (2019-2022), calculated using the *DynaRankR*-package (Strauss & Holekamp, 2019)

Dominance ranks were calculated using randomized elo-ratings, and all individuals with at least one interactions at a site were included in the analysis.

| Level | Time period | Roosting site | Similarity | Random expectation |
|---|---|---|---|---|
| All ages and sexes | July & September 2019 | BA | 0.66 [0.65 - 0.67] | 0.50 [0.46 - 0.54] |
| | July & September 2019 | CG | 0.81 [0.80 - 0.83] | 0.50 [0.44 - 0.56] |
| | July & September 2019 | NB | 0.71 [0.70 - 0.73] | 0.50 [0.43 - 0.56] |
| | 2019 & 2022 | CG | 0.81 [0.78 - 0.83] | 0.50 [0.43 - 0.56] |
| Juveniles | July & September 2019 | BA | 0.60 [0.56, 0.64] | 0.50 [0.39, 0.61] |
| | July & September 2019 | CG | 0.69 [0.66, 0.73] | 0.50 [0.39, 0.61] |
| | July & September 2019 | NB | / | / |
| | 2019 & 2022 | CG | / | / |
| Adults | July & September 2019 | BA | 0.72 [0.70 - 0.74] | 0.50 [0.42 - 0.58] |
| | July & September 2019 | CG | 0.78 [0.77 - 0.80] | 0.50 [0.44 - 0.56] |
| | July & September 2019 | NB | / | / |
| | 2019 & 2022 | CG | 0.82 [0.80 - 0.85] | 0.50 [0.43 - 0.57] |
| Females | July & September 2019 | BA | 0.63 [0.59 - 0.66] | 0.50 [0.40 - 0.60] |
| | July & September 2019 | CG | 0.60 [0.58 - 0.63] | 0.50 [0.43 - 0.57] |
| | July & September 2019 | NB | / | / |
| | 2019 & 2022 | CG | 0.87 [0.83 - 0.92] | 0.50 [0.38 - 0.63] |
| Males | July & September 2019 | BA | 0.66 [0.64 - 0.69] | 0.50 [0.43 - 0.57] |
| | July & September 2019 | CG | 0.85 [0.83 - 0.87] | 0.50 [0.42 - 0.58] |
| | July & September 2019 | NB | / | / |
| | 2019 & 2022 | CG | 0.67 [0.63 - 0.70] | 0.50 [0.41 - 0.59] |
